# SNP discovery by exome capture and resequencing in a pea genetic resource collection

**DOI:** 10.1101/2022.08.03.502586

**Authors:** G. Aubert, J. Kreplak, M. Leveugle, H. Duborjal, A. Klein, K. Boucherot, E. Vieille, M. Chabert-Martinello, C. Cruaud, V. Bourion, I. Lejeune-Hénaut, M.L. Pilet-Nayel, Y. Bouchenak-Khelladi, N. Francillonne, N. Tayeh, J.P. Pichon, N. Rivière, J. Burstin

## Abstract

In addition to being the model plant used by Mendel^1^ to establish genetic laws, pea (*Pisum sativum* L., 2n=14) is a major pulse crop cultivated in many temperate regions of the world. In order to face new challenges imposed particularly by global climate change and new regulations targeted at reducing chemical inputs, pea breeders have to take advantage of the genetic diversity present in the *Pisum* genepool to develop improved, resilient varieties. The aim of this study was to assess the genetic diversity of a pea germplasm collection and allow genome-wide association studies using this collection.

To be able to perform genome-wide association approaches with high resolution, genotyping with a large set of genetic markers such as Single Nucleotide Polymorphism (SNP) markers well-spread over the genome is required. Rapid advances in second-generation sequencing technologies and the development of bioinformatic tools have revolutionized the access to and the characterization of available genetic diversity. High-density, high-throughput genotyping has been possible for a large number of species, including those with large and complex genomes^2^ such as pea (2n=14) which genome size is estimated to be 4.45 Gb^3^. In this study, which is part of the PeaMUST project^4^, we used a target capture technology based on pea transcriptome sequences to generate exome-enriched genomic libraries that were further subjected to Illumina sequencing in paired-end mode. This methodology was chosen because whole-genome resequencing is relatively expensive for species with large genomes and because capturing genetic variations in repeated non-coding regions is difficult to achieve or to interpret^5^. Whole-exome sequencing represented an interesting alternative that focused on coding regions only^6,7^. Mapping the obtained reads on the reference pea genome sequence enabled the discovery of an abundant set of SNPs. The development of this resource is a crucial cornerstone in research and breeding projects towards boosting the improvement of pea production and quality.

## Methods

### Plant material and DNA extraction

A set of 240 *Pisum* accessions was selected, including 220 accessions originated from a larger panel structured into 16 genetic groups^8^ and 20 additional accessions chosen for their phenotypes. The 240-accession collection is referred to as Architecture and Multi-Stress (AMS) collection, since the accessions represent a broad diversity for root and shoot architecture and for biotic and abiotic stress responses. This collection contains cultivars, landraces, and wild types (including some *Pisum fulvum* and *Pisum sativum* subspecies accessions) with diverse geographical origins (Supplementary Table 1). Leaves were collected from 10 plants per accession, flash frozen in liquid nitrogen and stored at -80°C prior to DNA extraction. Tissues were then ground in liquid nitrogen using a pestle and a mortar. Genomic DNA extraction was performed using Nucleospin PlantII minikit (Macherey‐ Nagel, Hoerdt, France) following the manufacturer’s instructions.

### Probe design

As the pea genome sequence was not available at the time the probe design was made, two pea transcriptome datasets^9,10^ were used to build a reference set (refset) of expressed genes. After redundancy was removed, 67,161 unique contigs were kept, 20,972 of them being common to the two sequence datasets. The first exome capture design based on the refset was undertaken after predicting putative exon/intron junctions, masking repetitive sequences as well as excluding putative mitochondrial and chloroplastic sequences. A first probe design was performed by Roche™ (Madison, WI, USA) targeting 68.3 Mb. The analysis of the first capture results with the original probes demonstrated that a minority of 10-15% of the contigs retained the majority of the sequencing efforts, resulting in insufficient coverage for the remaining contigs. Sequencing data were used to identify the repetitive regions and a new probe design was performed after masking them to target a final capture space of 41.3Mb, representing 51,225 cDNA contigs.

### Library preparation and target enrichment

DNA samples were normalized before being fragmented with Adaptive Focused Acoustics® Technology (Covaris Inc., Massachusetts, USA). A 250-bp target size was obtained by using a Covaris E220 system, according to the manufacturer’s instructions. Then DNA fragments underwent a NGS library preparation procedure consisting in end repair and Illumina adaptor ligation using the KAPA HTP kit (Roche, Basel, Switzerland). Individual index sequences were added to each library for identifying reads and sorting them according to their initial origins. The Sequence Capture was performed using SeqCap EZ Developer kit^11^ according to the manufacturer’s instructions (Roche™). The sequence capture reaction efficiency was evaluated by measuring, using quantitative PCR, a relative fold enrichment and loss of respectively targeted and non-targeted regions before and after the sequence capture reaction.

### Targeted resequencing, sequence alignment and SNP calling

The captured samples were sequenced on HiSeq 2000 sequencing platform (Illumina, California, USA) with a PairedEnd sequencing strategy of 2 reads of 100 bases. The sequenced reads were trimmed for adaptor sequences using cutadapt 1.8.3^12^. Low-quality nucleotides with quality value<20 were removed from both ends. The longest sequence without adapters and low-quality bases was kept. Sequences between the second unknown nucleotide (N) and the end of the read were also trimmed. Reads shorter than 30 nucleotides after trimming were discarded. These trimming steps were achieved using fastx_clean (http://www.genoscope.cns.fr/fastxtend), an internal software based on the FASTX library (http://hannonlab.cshl.edu/fastx_toolkit/index.html). The reads were then aligned on the targeted regions with Novoalign V3.09 (http://www.novocraft.com, Selangor, Malaysia). We took advantage of the v1 genome sequence of cv. Cameor published meanwhile^13^ to perform SNP detection. Single nucleotide variants were detected on all the samples using samtools mpileup, followed by bcftools call and bcftools filter^14^ with a minimum genotype quality of 20 and a minimum coverage of 5 reads per sample. SNP variants for which more than 10 percent of the samples in the panel were heterozygous were then filtered out.

### Phylogenetic analysis

A subset of 206,474 SNPs was selected from the Dataset by applying filters on missing data (<20%), Minor Allele Frequency (>1%) and linkage disequilibrium (LD pruning based PLINK --indep 50 5 2). This subset was used to build a maximum likelihood phylogenetic tree using IQtree^15^, version 2.1.2, with model of substitution GTR+F+ASC. Alternatively, we also used a coalescent approach using 10,000 SNP non-overlapping windows as described by Wang et al^16^. The trees were also constructed using IQtree with the same parameters and finally were summarized using ASTRAL v5.15.1^17^.

Tree visualizations were generated using R package ggtree^18^.

### Structure analysis

Structure within the collection was calculated using the Bayesian clustering program FastStructure^19^ using a logistic prior for K ranging from 1 to 10. The script chooseK.py (part of the FastStructure distribution) was used to determine the best K that explained the structure in the collection based on model complexity. In addition, discriminant analysis of principal components (DAPC) was applied using DAPC function from adegenet package (version 2.1.5)^20^ in order to describe the genetic structure of the panel. Both FastStructure and DAPC groups were visualized on the phylogenetic tree using R package ggtreeExtra^18^.

### Data Records

The collection of 240 pea accessions was genotyped using an original set of probes for exome capture. Sequence data are available on NCBI (Bioproject PRJEB56612) and number of sequencing reads per accession are listed in Supplementary Table 2. Using the pea Cameor genomic sequence v1^13^ as a reference, 2,285,342 SNPs were identified. The full set of variants has been recorded as a VCF file and deposited at URGI (https://doi.org/10.15454/3QRIPA). The variant statistics per accession are reported in Supplementary Table 3. In average, 183,170 homozygous variants (compared to Cameor genome sequence) were detected per accession. As expected, the genotyping data from Cameor (DCG0251) used as an accession in the AMS panel were overall conform to the reference alleles from the reference genome sequence. Differences were only seen for 121 positions (0,005%). Among the detected SNPs, 647,220 were singletons (detected in only one accession). Excluding the reference Cameor (DCG0251), the number of singletons per accession ranged between 2 for VKL0176, a cultivated winter fodder pea to 113,086 for VSD0034, a *Pisum sativum* subsp. *abyssinicum* accession. The set of identified polymorphisms spans the seven pseudomolecules of the Cameor genome assembly at a frequency varying from 1 variant every 1831 bases for chromosome 2 to one every 1345 bases for chromosome 1 (Supplementary Table 4).

### Variant effects

We used the snpEff program^21^ in order to categorize the detected SNPs according to their predicted effects or their locations (Supplementary Table 4). The vast majority (76,71%) were labelled as “Modifier” (placed upstream or downstream of genes) and 14.32%, 8.76%, and 0.22% of the SNPs had a low (no change of the the protein sequence), moderate (change of aminoacid), and high (major change in the protein) predicted impact on gene functions, respectively. In fact, out of the total SNPs detected in coding regions (23.29%), 57.52% were predicted to be silent, 41.95% were predicted to induce an amino acid change in the coded protein and 0.53% were predicted to have a disruptive nonsense effect (premature stop codon, splicing junction modification or start codon missing).

Our data provide insights into exome genetic variation and highlight mutations with functional effects. This polymorphism inventory is valuable to explain the phenotypic diversity in the *Pisum* species.

### Phylogeny and structure analysis

Three different complementary approaches (Discriminant analysis of principal components (DAPC), FastSTRUCTURE, Maximum Likelihood Phylogenetic trees using both standard substitution model and a coalescence-based approach have been used to study the structure of the germplasm panel.

DAPC led to a clustering of the accessions in seven groups, as the most likely structure according to Bayesian information criterion. Clustering (Supplementary Table 1, Figure 1) tended to separate accessions according to crop evolution and cultivation types. Cluster 1 consisted in 14 accessions, mainly landraces or primitive germplasm. Cluster 2 comprised 87 accessions with an important proportion of spring cultivars including garden peas. Cluster 3 is composed of 27 accessions with a majority of spring-type lines coming from an Aphanomyces root rot breeding programme from Groupement des Sélectionneurs de Protéagineux (GSP, France) and Cluster 4 was a mix of spring-type and winter-type field peas with three primitive or landrace accessions. Cluster 5 grouped 5 wild-type accessions including *Pisum fulvum*, and Clusters 6 and 7 included winter-type field pea and fodder pea cultivars, respectively.

**Figure 1:**
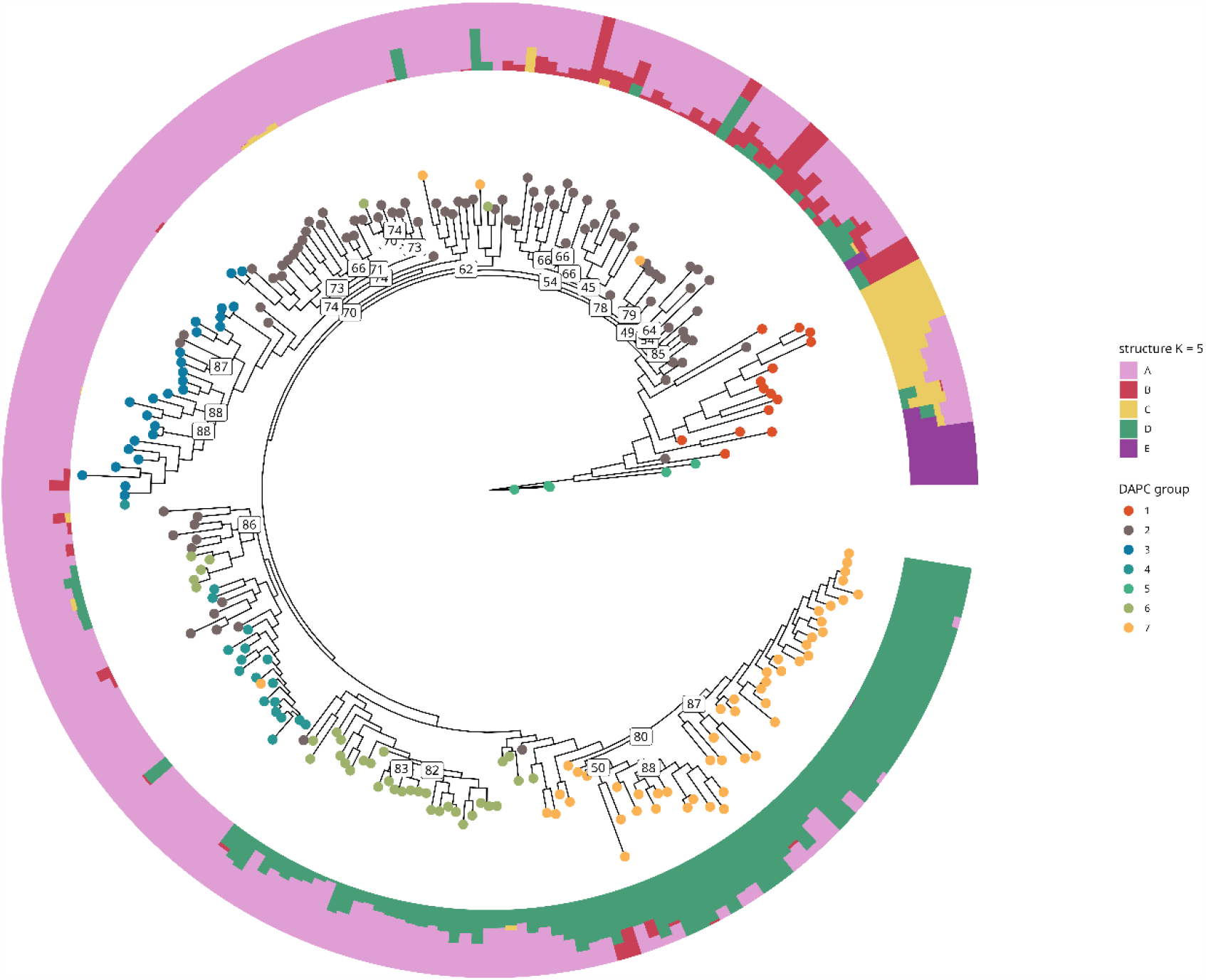
Maximum-likelihood phylogenetic analysis using 206,474 SNPs, DAPC grouping, and FastStructure composition of the 240-accession pea AMS collection. Colour codes indicate the different groups as inferred by DAPC and FastStructure. Numbers shown at the nodes show bootstrap support when below 90%.

FastStructure, on the other hand, inferred that the panel could be divided into five ancestral subpopulations (numbered A to E, Figure 1). Subpopulation E corresponded to the DAPC cluster of 5 wild accessions with very little admixture with the other clusters while subpopulation C corresponded to Cluster 1 (landraces/primitive germplasm). Subpopulation D corresponded to the fodder pea cluster 7. Clusters 3,4 and 6 (field pea cultivars mainly) seemed to derive from the ancestral subpopulation A. Some admixture between subpopulations A and D was observed for the winter field pea accessions from Cluster 6, and between subpopulations A, B and D for some garden pea cultivars.

The Phylogenetic tree (Figure 1, produced using the 206,474 SNPs) confirmed the DAPC and FastStructure observations with only few placement differences. The summarized phylogenetic tree inferred with the coalescent approach (Supplementary Figure 1) corroborated the clade delimitations. However, the relationships between clades differed (Supplementary Figure 2) which could be attributed to admixture during cultivar selection leading to incomplete lineage sorting.

In conclusion, this dataset is a large marker resource that can be used for different purposes, including the development of targeted genotyping tools for molecular identification, genetic mapping or genomic selection in pea. It provides insights into pea diversity and helps to investigate selection processes in this species. The SNP resource also empowers Genome-wide Association Studies targeted at revealing the genetic architecture of important traits and highlighting alleles to be used in pea breeding programmes. Indeed, the collection has been evaluated for different traits including plant architecture, phenology and resistance or tolerance to a range of biotic and abiotic stresses, as exemplified by Ollivier et al^22^ who deciphered the genetic determinism of resistance to two aphid biotypes.

## Acknowledgements

This study is part of the PeaMUST project, that was funded by the French government through the Investment for the Future program (project ANR-11-BTBR-0002) and Ministère de l’Enseignement supérieur, de la Recherche et de l’Innovation.

## Author Contributions

ILH, MLPN, VB and JB designed the diversity panel. Experiments were conceived and designed by GA, NR, JPP and JB, and performed by HD, AK, KB, EV, MCM and CC. Data were analysed by ML, JK, YBK and GA and NF organised the data access. The manuscript was written by GA and JK and all authors read and improved it.

