## Supplementary Figures 1-2 for "SNP discovery by exome capture and resequencing in a pea genetic resource collection"

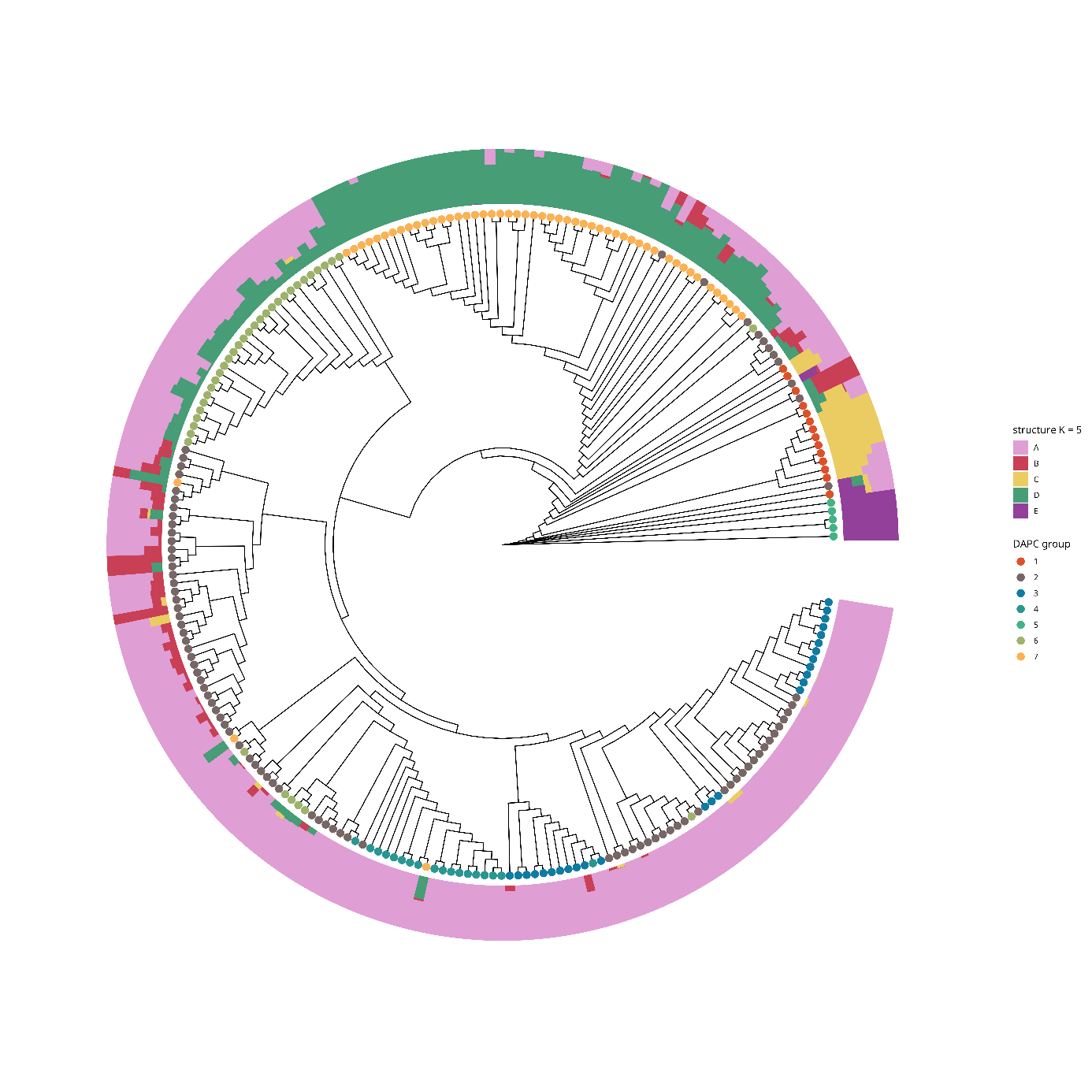

**Supplementary Figure 1:** Summarized Phylogenetic tree obtained with a coalescent approach using 10,000 SNPs non-overlapping windows. DAPC grouping, and FastStructure are indicated for each of the 240-accession pea AMS collection. Colour codes indicate the different groups as inferred by DAPC and FastStructure.

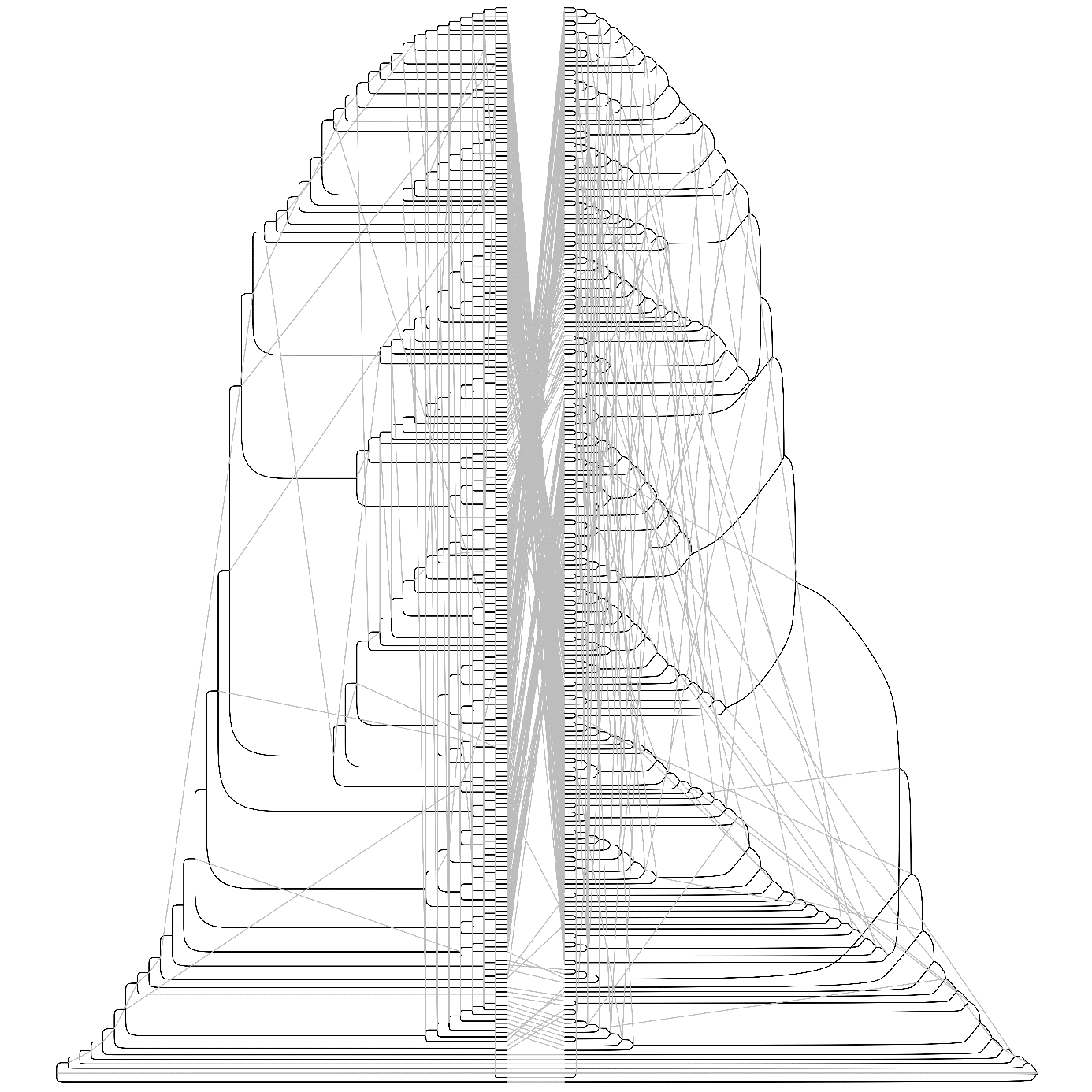

**Supplementary Figure 2:** Comparison of the phylogenetic tree obtained in Figure 1 (left) and the summarized tree inferred with the coalescent approach (right). Figure was generated using GGTree
